# Whole genome analyses of the endangered Northern abalone (*Haliotis kamtschatkana*) reveal population differentiation and a genomic signature of a dramatic population decline

**DOI:** 10.1101/2025.09.17.676880

**Authors:** Anna Tigano, Erin C. Herder, Kayla Long, Janine Supernault, Daniel L. Curtis, S. Christine Hansen, Mackenzie D. Mazur, Eric B. Rondeau, Dominique Bureau

## Abstract

Despite widespread declines of many wildlife species, the effects of population decline on the genetic health and the recovery potential of affected species is still poorly understood, especially beyond a few charismatic species. The Northern abalone (or Pinto abalone; *Haliotis kamtschatkana*) is a marine gastropod mollusc of social, cultural and historical economic importance in the Pacific Northwest of North America that experienced a decline in population density due to commercial harvest and is currently listed as endangered in Canada under the Species at Risk Act. Previous genetic investigations based on microsatellites and reduced-representation approaches concluded that Northern abalone is panmictic throughout its range, from Alaska to California, and identified high levels of genetic variation with no indication of population decline. Using whole genome resequencing data from Northern abalone sampled across the northern part of the species range, we instead identified both: 1) significant differentiation between two genetic groups, albeit very concentrated in the genome; and 2) a strong signature of a dramatic population decline, without evidence of genetic inbreeding. Even though demographic reconstructions showed a timid signal of recent population expansion, the pervasive excess of rare alleles identified (including a high occurrence of singletons) may pose a genetic load risk, potentially hindering the species recovery. We also found evidence of historical, rather than current, connectivity throughout the area investigated. These results are important for management decisions and highlight the utility of whole genome data in conservation, especially in species with historically large effective population sizes like the Northern abalone.

## Introduction

Species losses and population declines are occurring at alarming rates and are mostly attributed to the direct and cascading effects of anthropogenic activities (Cowie et al., 2022). While severe population declines have been reported in thousands of species (WWF 2024), the effects of population bottlenecks on the genetic health, and the ability to recover, of these declining species are not well understood, and are hence difficult to predict. For example, whole genome analyses have shown low inbreeding load and no signs of inbreeding depression in the critically endangered vaquita porpoise (*Phocoena sinus*), even though only ∼10 individuals remain in the wild (Robinson et al., 2022). In contrast, the endangered Southern resident killer whale population (*Orcinus orca*) is more abundant than the vaquita with ∼70 individuals, yet genomic analyses have revealed substantial inbreeding depression and resultant health issues that are preventing the population’s recovery (Kardos et al., 2023a). In fact, while the vaquita is thought to have some recovery potential if bycatch mortality is reduced or eliminated, the southern resident killer whale has not shown signs of recovery despite ∼50 years of conservation efforts (Kardos et al., 2023; Robinson et al., 2022). The discovery of contrasting genomic signatures of population decline in these two relatively closely related species highlights the usefulness of genomic data in making links between bottleneck and inbreeding depression, and characterizing extinction risks.

Insights into inbreeding depression and extinction risks at the genomic level in the vaquita and killer whale examples above were obtained from the analysis of whole genome data, which have greatly contributed to the field of conservation genomics (Supple & Shapiro, 2018; Taylor et al., 2021). The analysis of whole genome data enables the complete characterization of levels and distribution of genomic variation across the genome, which provides information on inbreeding depression, adaptive potential, and potential for population collapse or recovery. Furthermore, whole genome data enable the reconstruction of demographic histories. Considered together, this information is crucial to characterize the genetic health of populations and inform their management and conservation. For example, healthy levels of genetic diversity reduce the risk of genetic load and represent a proxy for the potential of populations and species to adapt to changing conditions in their environment, which enables their long-term persistence (Kardos et al., 2021). Reconstruction of demographic histories can elucidate the factors leading to population declines, validate empirical historical observations and reconstructions from other sources, or provide a reference baseline in their absence (Hohenlohe et al., 2021). Additionally, whole genome analyses can reveal genomic differentiation that can potentially go undetected with the analysis of microsatellites or reduced-representation approaches such as RAD-seq. The increased resolution granted by the full coverage of the genome can result in a better understanding of population structure and its drivers (St. John et al., 2025) and in the identification of markers able to discern populations and species (Toews et al., 2016).

While a few vertebrate species have been intensely studied to investigate the genomic signature of their decline (e.g., Kardos et al., 2023b; Khan et al., 2021; Robinson et al., 2022), invertebrates-representing 95% of animal biodiversity-are heavily underrepresented in conservation genomics research (Lopez et al. 2019). Given their dramatically different and diverse life history traits, a better understanding of invertebrate genomic health, especially in declining and endangered species, is needed. Characterizing patterns and drivers of population differentiation could help identify management strategies and conservation priorities, aid in the design of protected areas, and provide insights into the variety of biotic and abiotic factors affecting genomic health and extinction risks across species. Abalone, for example, are large marine gastropod molluscs with more than 50 species worldwide and are known for their economic importance as a seafood delicacy. Many abalone species have suffered population declines driven by overharvesting and illegal harvest (Cook and Shumway, 2023), and some species have suffered high mortality from diseases (Rogers-Bennett et al., 2002). The Northern abalone (or Pinto abalone; *Haliotis kamtschatkana*) has a wide distribution, from Southeast (SE) Alaska (USA) to Baja California (Mexico). The Northern abalone experienced a dramatic population decline across its range, and is currently listed as Endangered in Canada (Species at Risk Act (SARA) Species Registry) and on the International Union for Conservation of Nature (IUCN) Red List (Peters and Rogers-Bennett, 2021). Overharvesting was the major driver of population decline in Canada and all commercial, recreational and First Nations’ food, social and ceremonial fisheries were closed in 1990 (Campbell, 2000). Nonetheless, illegal harvesting continues to be a threat for the species (Neuman et al., 2018).

Despite the well-documented decline of Northern abalone, previous studies based on 12 microsatellites (Withler et al. 2003) and ∼6,000 SNPs (Dimond et al., 2024) found overall high levels of genetic variation, and estimated large effective population sizes with no signature of population bottleneck. Furthermore, both studies, covering either the northern part of the species range, i.e. British Columbia (BC) and Southeastern (SE) Alaska (Withler et al., 2003), or the entire species range (Dimond et al., 2024), did not identify significant population structure and differentiation, concluding that the species is panmictic across its whole range from Mexico to Alaska. Lack of population structure was attributed to levels of current or historical gene flow sufficient to homogenize genetic variation across a large geographical range. However, panmixia is difficult to reconcile with the species’ patchy distribution (spanning over 3,700 km), and life history traits, including a relatively short planktonic phase (7-10 days; Strathmann 1987) and low adult dispersal (Sloan and Breen 1988). As a comparison, marine invertebrates with longer planktonic phases, such as the bat star (*Patiria miniata*; 6-10 weeks) and the giant red sea cucumber (*Apostichopus californicus*; 2-4 months) have shown significant population structure within similar ranges as the Northern abalone, even with fewer markers (7 microsatellites and ∼3,000 SNPs, respectively; Sunday et al., 2014; Xuereb et al., 2018). It is therefore plausible that the genetic markers previously used to investigate the genetic health and population structure of Northern abalone were not sufficient to identify signatures of differentiation and population decline due to limited resolution, high levels of genetic variation, and/or concentrated genomic signatures of bottleneck and differentiation.

In Canada, Northern abalone is managed on a coastwide basis as a single Designatable Unit (DU; COSEWIC 2009) based on inferences about species genetic discreteness and uniqueness from a reduced representation of the genome. Here, we used whole genome data from 15 locations across the northern part of the Northern abalone range (BC and SE Alaska) to: 1) characterize population structure and differentiation, if any; 2) determine the effect of the severe species decline in the genome; and 3) explore the relative roles of the factors that either hinder or explain good genomic health and panmixia despite expectations. While whole genome data may corroborate a lack of population structure and genomic variation indicative of good genetic health, analysing genomic variation and differentiation across the whole genome will grant maximum resolution and definitive answers. The results from this work will provide important information for the management of Northern abalone in Canada and a better understanding of the effect of population declines in understudied taxa such as molluscs.

## Methods

### Library preparation and sequencing

We used epipodial tissue samples that were collected non-destructively between 1999-2019 from 111 individuals from 15 sampling locations; 14 locations along the BC coast and one location in SE Alaska, as per the methods described in Withler et al. (2003) and a series of SARA collection permits for the work (Table 1; Figure 1). We extracted DNA using a Qiagen Biosprint 96 DNA extraction kit. Whole genome library preparation and sequencing was performed at Genome Québec. Genomic DNA was quantified using the Quant-iT™ PicoGreen® dsDNA Assay Kit (Life Technologies). Libraries were generated from 100 ng of DNA using the NEBNext Ultra II DNA Library Prep Kit for Illumina (New England BioLabs) as per the manufacturer’s recommendations. Size selection of libraries was performed using SparQ beads (Qiagen). Libraries were quantified using the KAPA Library Quantification Kits-Complete kit (Universal; Kapa Biosystems) and their average fragment size was determined using a Fragment Analyzer 5300 (Agilent) instrument. The libraries were normalized, pooled, and sequenced on an Illumina NovaSeq X platform using 150 bp paired-end reads.

**Figure 1.**
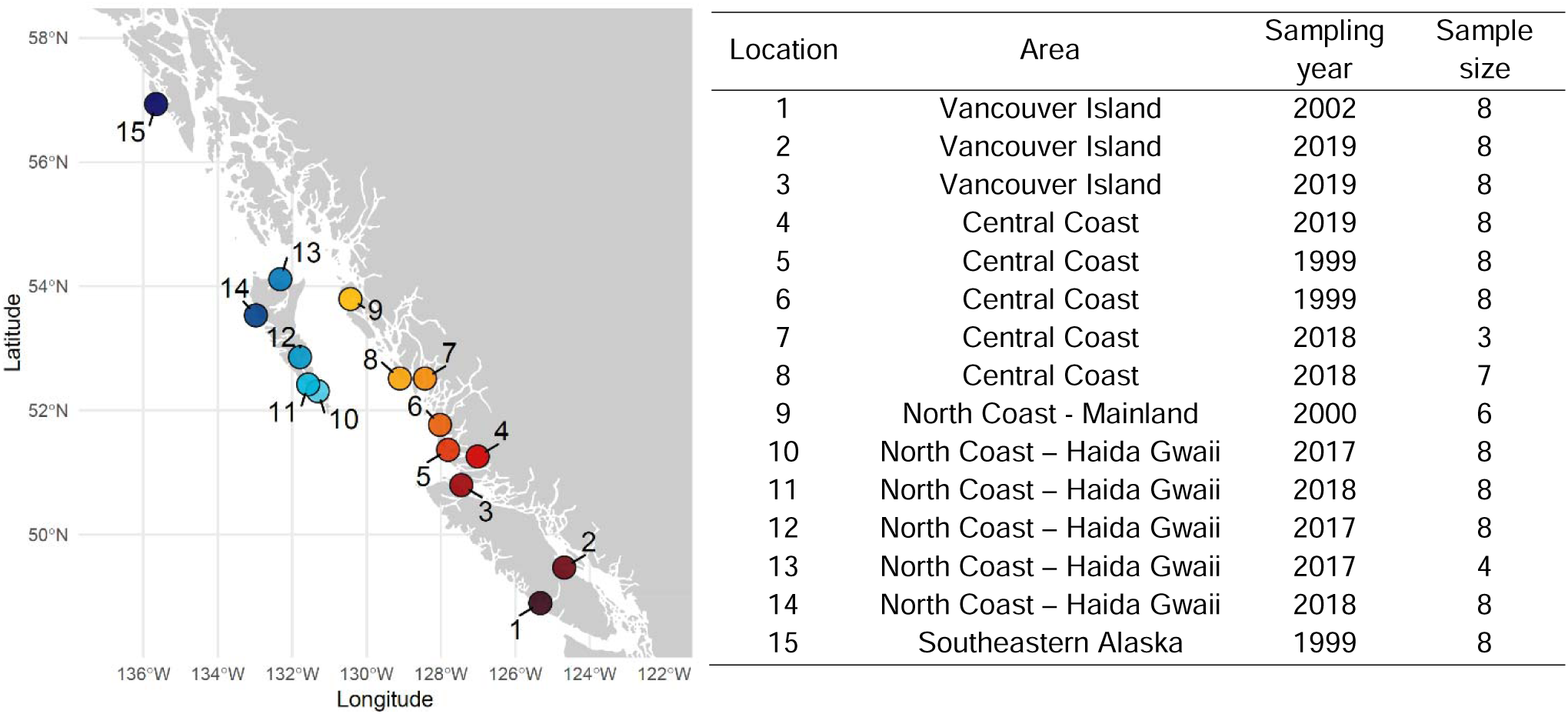
Map showing sampling locations and table summarizing area and sample size information. Coordinates and precise locations are restricted for the endangered Northern abalone under Section 124 of the Species at Risk Act.

### Variant calling

We used the snpArcher pipeline for variant calling (Mirchandani et al., 2024). SnpArcher takes raw sequencing data, processes them, and calls sequence variants from whole genome resequencing data. We used GATK (McKenna et al., 2010; Van der Auwera et al., 2013) as the variant caller, with all default settings other than selecting the low coverage option. We ran the pipeline twice using two different reference genomes for read mapping. In the absence of a reference genome for the Northern abalone, we used the black abalone (*Haliotis cracherodii*) reference genome as it is the most contiguous abalone reference genome available. We then repeated variant calling using the red abalone (*Haliotis rufescens*) reference genome, as it is more closely related to Northern abalone and is gene annotated. The proportion of mapped reads (98.6% versus 99.1%), the proportion of properly paired reads (87.7% versus 91.7%), and the number of high-quality SNPs (47 million SNPs versus 56 million SNPs, see Results) were higher when the data were mapped to the red abalone genome compared to the black abalone genome, so we retained the red abalone reference genome for subsequent analyses. Note that we renamed reference genome scaffolds with numbers in order from largest to smallest.

We applied additional quality filters to the raw variants output by snpArcher with VCFtools v0.1.16 (Danecek et al., 2011). We filtered out variants with depth 4 and 30 and with more than 30% missing data, kept only biallelic sites and removed indels (“full dataset”). We then split the variants from the full dataset into two additional datasets based on their minor allele frequency (MAF) calculated across all individuals: 1) the ‘common alleles’ dataset included variants with MAF ≥ 0.05 and 2) the ‘rare alleles’ dataset including variants with MAF < 0.05. The three datasets (full, common, and rare) were used for different analyses downstream: the full dataset was used for unbiased measures of genetic diversity and demographic reconstructions, the common allele dataset was used for estimates of genomic differentiation between locations and groups, and the rare alleles dataset was used to test hypotheses about population connectivity and demography.

Preliminary analyses, including a Principal Component Analysis (PCA), and coverage analysis, identified two individuals showing anomalous divergence from the rest of individuals probably due to sample contamination, and one individual with more than 30% missing data due to low genome coverage. These three individuals were excluded from downstream analyses (n=108).

### Population structure and patterns of genomic differentiation

We explored population structure using a PCA with the package SNPRelate (Zheng et al., 2012) in R 4.4.1 (R Core Team, 2024). We first used a random subset of 100,000 common allele SNPs to conduct an exploratory PCA. We then tested for the minimum number of SNPs necessary to retrieve a signal of population structure by performing PCAs with 10,000, 1,000, 100 and 10 MAF-filtered SNPs. We also ran a PCA with 1 million SNPs to test whether increasing the number of SNPs would reveal finer population structure.

From these analyses we identified two genetic groups: a southeast group, including locations 1-9, and a northwest group, including locations 10-15 (see Results). To characterise patterns of differentiation in the genome and identify differentiated loci between these two genetic groups, we estimated F_ST_ at each individual SNP and in 50-kb moving windows using VCFtools. Then, we selected the most differentiated SNPs, i.e. the SNPs showing the highest F_ST_ between the southeast and the northwest groups, created subsets including the 10, 100, 1,000, 10,000, 100,000 and 1 million most differentiated SNPs respectively, and performed PCAs using each of these sets of SNPs.

We further examined the 1,000 most differentiated SNPs in the genome. We calculated their MAF in each location and across groups, and their difference in allele frequencies (ΔAF) between the northwest and the southeast groups. Furthermore, we considered these 1,000 SNPs for further functional exploration. We functionally annotated the SNPs including the genes that were directly affected (i.e. the SNP that fell on the gene sequence) and the genes that were closest to SNPs falling in intergenic areas (based on the assumption that SNPs in intergenic areas, potentially regulatory, would affect the closest gene in the linear genome) using BEDtools functions in the *valr* v0.8.4 package in R (Quinlan & Hall, 2010; Riemondy et al., 2017). We used various databases, including SwissProt, NCBI, and GeneCards, (Boutet et al., 2007; NCBI 1988; Stelzer et al., 2016) to gather information on the function of genes associated with differentiated variants.

### Genomic diversity

To assess levels and distribution of sequence variation among samples and across the genome, we calculated nucleotide diversity and Tajima’s D in 50-kb moving windows using the full dataset across all individuals combined, and separately across each of the two identified genetic groups (northwestern and the southeastern groups, see Results). We also calculated individual heterozygosity using the full and the common alleles dataset. All analyses were performed in VCFtools.

### Demographic reconstructions

We reconstructed the demographic history of Northern abalone with GONE (Santiago et al., 2020), a software that uses patterns of Linkage Disequilibrium (LD) between SNPs to estimate changes in effective population size (*N_e_*) up to 140 generations in the past, where generation is defined as the average age of reproducing individuals in a population. We set a maximum of 50,000 SNPs per chromosome, a recombination rate of 1 cM/Mb (representing the mean recombination rate in molluscs (Stapley et al., 2017) in the absence of species-specific information), the Haldane correction for genetic distance, and 40 replicates. To generate the input files for these analyses, we thinned the full dataset, including all individuals, only northwest individuals, or only southeast individuals, respectively, and retained only SNPs that were located farther than 1 kb from each other and on scaffolds longer than 10 Mb, and then further randomly selected 500,000 SNPs. We repeated the demographic reconstructions using two additional recombination rate estimates, 0.5 and 2 cM/Mb, with the rest of the parameters kept the same, to explore the effect of varying recombination rate on the demographic reconstructions.

### Runs of homozygosity

To evaluate whether the recent population decline of Northern abalone has led to genetic inbreeding, we calculated Runs of Homozygosity (ROHs) in each individual using Plink v1.9 (Clarke et al., 2011). ROHs are long stretches of identical-by-descent (IBD) haplotypes that appear in individual genomes as the result of recent parental relatedness, and as such are unlikely to occur independently in the same position in different locations. We collapsed ROH segments identified across multiple individuals into unique ROHs if they overlapped by more than 50% and evaluated how many individuals and locations shared the same ROH segments. Based on their low homoplasy, we calculated the number of individuals sharing ROHs between locations with the *crossprod* function in R as an indicator of connectivity among sampling locations. As ROHs shorten with time due to recombination with non-homozygous traits, we can estimate ROHs coalescent times from their lengths (Foote et al., 2021; Thompson, 2013) and infer historical demographic events at varying points in time, including bottlenecks and historical connectivity. To estimate the age of ROHs in number of generations, we used the formula t = [100/(2 x L x r)] cM where t is coalescent time in number of generations, L is length in Mb, and r is recombination rate. In the absence of recombination rate estimates for abalone, we used the average recombination rate estimated for molluscs (1 cM/Mb; Stapley et al. 2017).

### Analysis of rare alleles

As an additional measure of connectivity and signature of population contraction and expansion, we examined the distribution of rare alleles within individuals and locations. We calculated the number of singletons and doubletons, i.e. the number of variants present in only one individual in either heterozygous or homozygous state (i.e. one or both alleles), and the number of locations each rare variant was shared across using VCFtools and R.

## Results

### Genomic diversity

We identified a total of 56.4 million high quality SNPs (full dataset), indicating, with 1 SNP every ∼24 bp on average, high overall genomic diversity. Of these SNPs, only 19.1 million (34%) had MAF > 0.05 (common alleles dataset). After quality filtering (including for MAF), high quality SNPs had an average sequencing depth of 9.7x (SD = 1.3x) and 9% (SD=3.6%) missing data.

Mean heterozygosity, i.e. the proportion of heterozygous sites in each individual, was 11.5% in the full dataset and 28.8% in the common allele dataset, with significant differences only between location 12, showing highest overall heterozygosity, and five other locations (1, 3, 4, 11, and 15), showing the lowest mean heterozygosity (Tukey Post-Hoc test, p <0.05). Mean nucleotide diversity across 50-kb windows was 4.17*10^-3^ across all individuals (n=108), 4.14*10^-3^ in the northwest (n=44), and 4.18*10^-3^ in the southeast (n=64). Mean Tajima’s D across all genomic 50-kb windows was negative across all individuals (Tajima’s D =-1.04), and within the southeast (Tajima’s D =-0.93) and the northwest group (Tajima’s D =-0.90; Supplementary Figure 1), indicating an overall excess of rare alleles.

### Population structure and genomic differentiation

The PCA based on 100,000 random SNPs revealed differentiation between two geographic groups: 1) Vancouver Island and the mainland coast of BC (locations 1-9), hereafter referred to as “Southeast group”, and 2) Haida Gwaii and SE Alaska (locations 10-15) termed the “Northwest group” (Figure 2C). Individuals were separated mostly along PC1, except one individual from site 13 in the Northwest group, clustering with individuals from the Southeast group. The PCAs based on less than 100,000 random SNPs showed decreasing group definition: with 10,000 SNPs, individuals tended to cluster within each of the two groups but did not form clearly distinct groups anymore, and with smaller datasets, individuals lost any population structure signal (Figure 2A-B). The PCA based on one million SNPs showed a similar clustering pattern as with 100,000 SNPs, with slightly stronger separation between the northwest and the southeast groups, and between location 15 (SE Alaska) and the rest of the northwest group individuals from Haida Gwaii (locations 10-14; Figure 2D).

**Figure 2.**
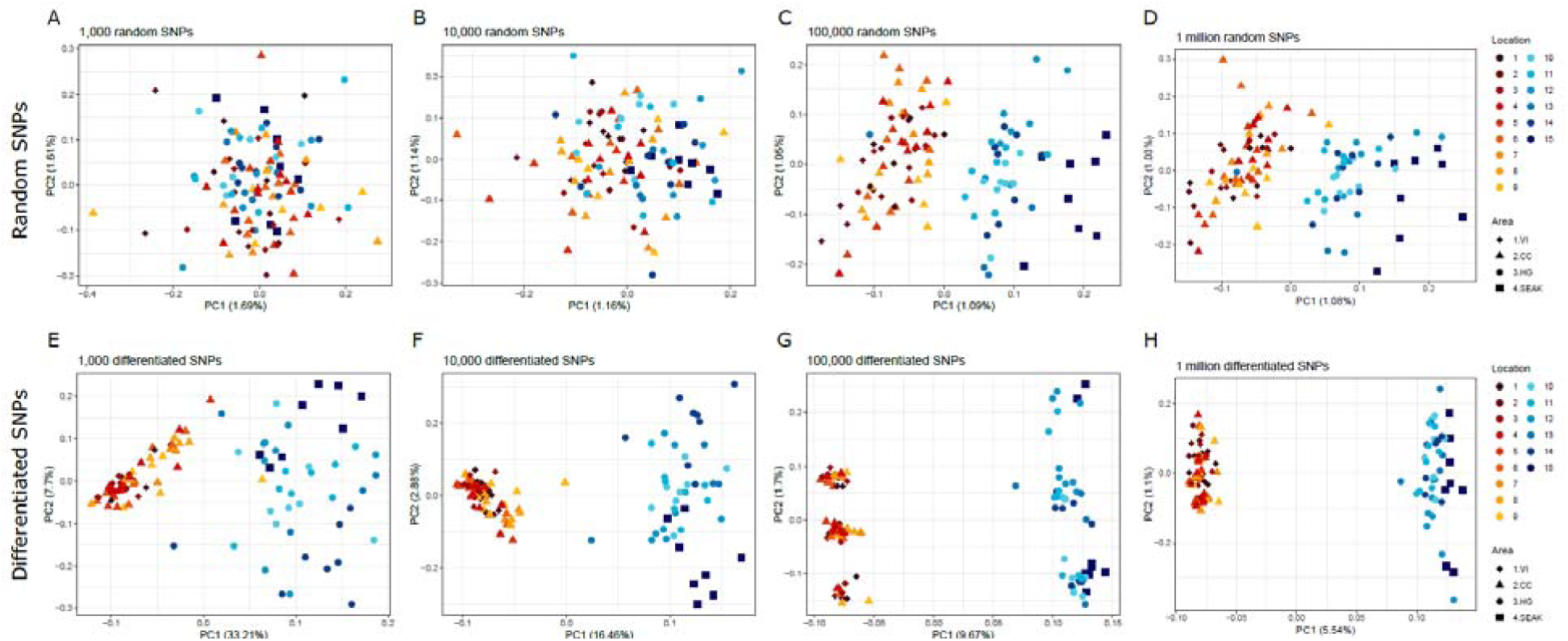
Plots of Principal Component Analyses based on an increasing number of random SNPs (A-D) and differentiated SNPs (E-H) from 1,000 on the left to 1 million on the right. VI = Vancouver Island, CC = Central Coast, HG = Haida Gwaii, and SEAK = Southeast Alaska.

The genomic landscape of differentiation between the two genetic groups showed overall low background F_ST_ (average F_ST_ across SNPs=0.001146 and across windows=0.001386) with a few 50-kb windows showing higher differentiation (Figure 3A). Genomic differentiation tended to be concentrated in a few regions of the genome: of the 27 windows with F_ST_ values above the 99.9^th^ percentile (“outlier windows”), eight were found on scaffold 9 and five on scaffold 4. Excluding three short scaffolds (< 125 kb) where outlier windows were likely artefacts, only seven additional scaffolds (for a total of nine) harboured outlier windows. Similarly, at the SNP level, of the 1,000 most differentiated SNPs, 404 were located on scaffold 9 and 223 were located on scaffold 4 (Figure 3B-C). The remainder were distributed across 22 additional scaffolds. However, only 12 of these scaffolds included more than 10 highly differentiated SNPs.

**Figure 3.**
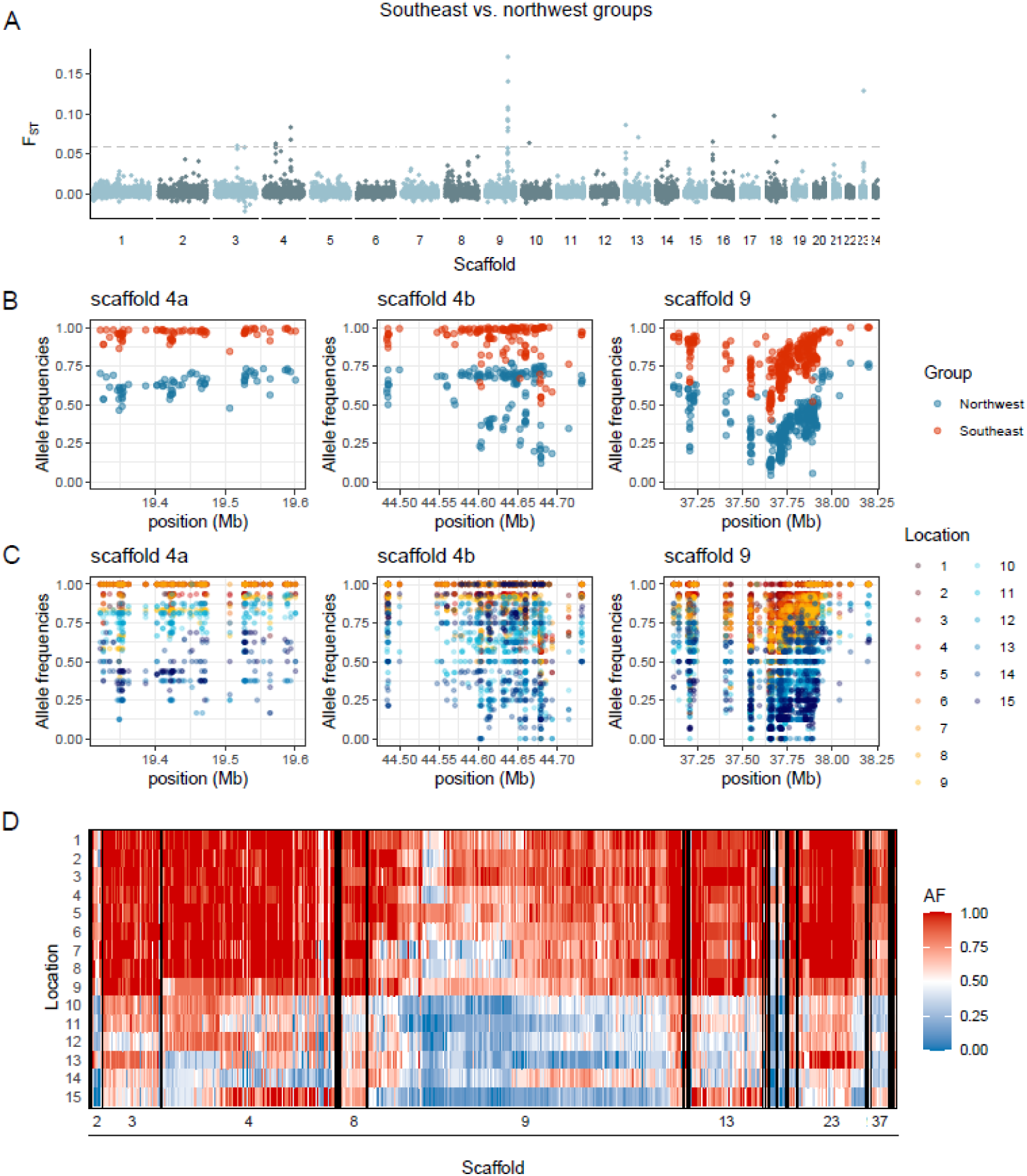
Patterns of genomic differentiation between the northwest and southeast groups. A) Manhattan plot showing patterns of differentiation calculated as F_ST_ in 50 kb moving windows across the genome. B-C) Plots showing variation in allele frequencies at each of the 1000 most differentiated SNPs located along the three main AODs on scaffold 4 (4a and 4b) and 9 calculated within genetic groups (B) and individual locations (C). D) Genotype plot summarizing allele frequency at each of the 1000 most differentiated SNPs across all scaffolds. The differentiated SNPs were located across 24 scaffolds, but only the eight scaffolds with the highest number of differentiated SNPs were labelled in the plot.

PCAs based on highly differentiated SNPs revealed a clear signature of population structure (Figure 2E-H), even with as little as the 10 most differentiated SNPs (Supplementary Figure 2). The PCAs based on the 1,000 and 10,000 most differentiated SNPs (Figure 2E-F) showed similar patterns, including a general higher variation in the northwest groups than in the southeast group. Using 100,000 and 1 million SNPs, individuals from the two groups started to appear strongly separated along PC1, with the only difference between the two PCAs being an additional grouping pattern along PC2 in the 100,000 SNP PCA, apparently unrelated to population of origin (Figure 2G-H).

None of the 1,000 most differentiated SNPs were fixed between the northwest and the southeast groups, with the maximum difference in allele frequency being 0.66 (Figure 3B-D). No SNP was fixed in the northwest group and variable in the southeast, but 65 SNPs were fixed in the southeast group and variable in the northwest, with 25 of these SNPs being fixed for the same allele at location 15, the SE Alaska location, as in the southeast group. Furthermore, 191-451 of the 1,000 most differentiated SNPs were fixed in each of the locations from the southeast group, while only 104 SNPs or fewer were fixed in each of the locations in the northwest group, again indicating higher variation in the latter. Of the SNPs that were fixed in more than one location in the southeast, only 0-4 were fixed in locations in the northwest for the same allele, except in the two northernmost locations, 13 and 15, which shared 54 and 81 fixed differences with locations from the southeast group, 50-81 more than the other northwest locations. Note that the fixed SNPs shared with the southeast were located in different parts of the genome in the two locations: those fixed in location 13 were mostly concentrated on scaffold 23, while those fixed in location 15 were mostly concentrated on scaffold 4 and 13, with no shared fixed differences between locations 13 and 15.

We identified 91 genes associated with the 1,000 most differentiated SNPs, 26 of which were uncharacterized (Supplementary Table 1). The two areas of differentiation (AOD) on scaffold 4 were associated with four and six genes, respectively. In the first AOD, only one gene was characterized, a zinc finger protein gene, while on the second AOD, the six annotated genes were mostly associated with immune and reproductive functions. The area of differentiation on scaffold 9 was associated with 14 genes, and eight of these were functionally annotated with functions mostly associated with immune functions, stress response, and apoptosis (Supplementary Table 1).

### Effective population size and population demography

To reconstruct the recent demographic history of Northern abalone in BC and SE Alaska we used multiple independent sources of information: 1) LD, 2) Tajima’s D, and 3) analysis of rare alleles. We first reconstructed the demographic history over time using genomic patterns of LD among SNPs based on the average mollusc recombination rate of 1 cM/Mb (Figure 4). Between 140 and 100 generations before present, both southeast and northwest groups appeared to have increased 2-3 fold their effective population size (*N_e_*) in 30-35 generations, which was followed by a long period of relative population stability. More recently, large declines in *N_e_* across BC and SE Alaska were observed. The northwest and southeast groups both showed a severe bottleneck, but differed in the timing of the population decline and their historical effective population sizes. Before the bottleneck, *N_e_* was relatively stable around 300,000 individuals in the southeast group, and around 500,000 in the northwest group. Effective population sizes started slowly declining in both groups approximately 39 generations before present, but the population bottlenecks occurred earlier in the southeast group. Northern abalone declined by more than 50% (compared to generation 39) at generation 25 before present in the southeast group and at generation 16 before present in the northwest group. From this point, *N_e_* in both groups further decreased to 10% in only 3-4 generations. *N_e_* reached its historical minimum 8-10 generations before present with a decline to 0.76% and 1.8% of their pre-bottleneck average in the northwest and in the southeast groups, respectively.

**Figure 4.**
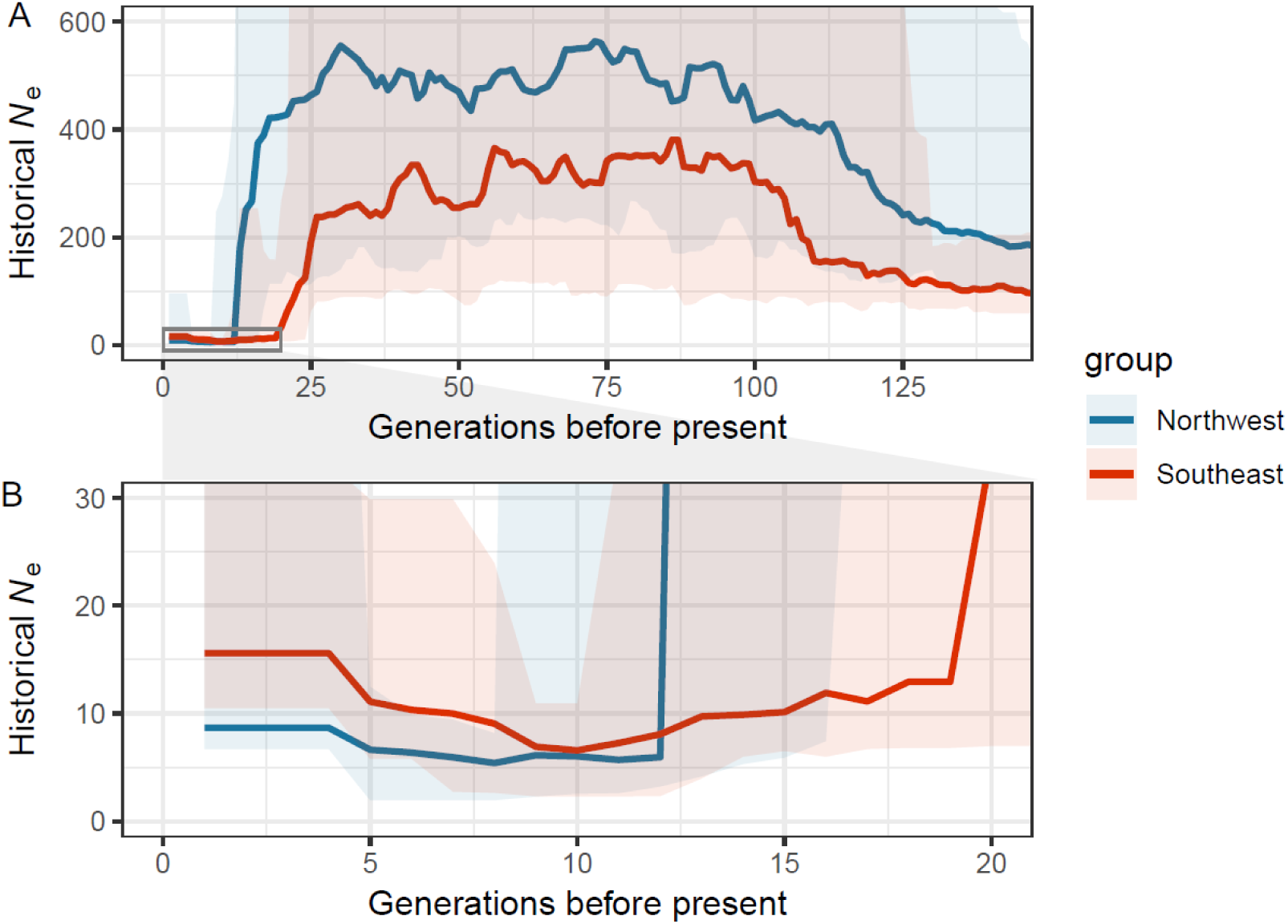
Plots summarizing the results from the demographic reconstructions from GONE A) for the last 140 generations and B) a zoom-in on the last 20 generations before sampling began in 1999. The thick solid line represents the median across 40 iterations and the shaded area the 95% confidence intervals.

GONE also detected a signature of recent population expansion in the last few generations, showing that *N_e_* almost tripled since the lowest estimate was reached approximately 8-10 generations before present in both groups. However, current *N_e_* estimates were only 5% and 2% of the average estimates from the 100 generations prior the bottleneck, in the southeast and northwest groups respectively.

The demographic reconstructions based on the two additional recombination rates (0.5 and 2 cM/Mb) detected the same overall patterns typical of a population bottleneck, but with expected differences in the estimated effective population sizes and timing of the bottlenecks. With r=0.5 cM/Mb, *N_e_* estimates were larger (up to 3 million individuals before the bottleneck in the northwestern group) and the bottleneck was dated further in time, at ∼30 and 50 generations before present in the northwestern and southeastern groups, respectively. In contrast, with r=2 cM/Mb *N_e_* estimates were smaller (reaching a maximum of 134,000 before the bottleneck in the northwestern group), the bottleneck occurred 5 generations before present, and no recent population expansion was detected (Supplementary Figure 3).

Estimates of Tajima’s D were negative across most of the genome, indicating an overall excess of rare alleles (Supplementary Figure 1), which being so pervasive across the genome, was indicative of population expansion rather than selection. Rare variants, those with overall minor allele frequency < 0.05, were abundant, making up 66% of all variants. The high proportion of rare variants found in only one individual or location (42%, and 28% of all variants) further suggests that a large number of new mutations arose recently, more so in the northwest group than the southeast group, based on the higher number of singletons in locations from the northwest (Supplementary Figure 4). These data indicate that these rare variants have not had the time to increase in frequency and to spread within and among locations, further supporting that the Northern abalone is slowly recovering from a relatively recent dramatic population bottleneck.

### ROHs calling

We identified a total of 88 ROH segments across 108 individuals. On average, individuals carried 3.4 ROHs (number of ROHs, NROH), covering 647 kb (sum of ROH lengths, SROH), or 0.05% of their total genome length, and ROH segments were 193 kb long on average. We observed variation among individuals, with eight individuals not carrying any ROHs, and two individuals carrying more than 10 ROHs. F_ROH_, a measure of inbreeding, is generally reported as the proportion of the genome affected by ROHs longer than 1.5 Mb, as a link has been suggested between ROHs of this length and inbreeding (McQuillan et al. 2008). In our dataset, the longest segment was 1.05 Mb, below the 1.5 Mb threshold. Of the 88 ROHs identified, 37 were found in one individual only and the rest of the segments were shared among two or more individuals and across multiple locations, with the exception of one segment on scaffold 14 that was identified in two individuals from the same location only. No ROH was private to any locations (Figure 5). Location 12 had the highest number of ROHs (n=41) while location 4 had the lowest number of ROHs (n=12) among the locations with a sample size of eight individuals. We did not observe any geographical patterns in the distribution of ROHs across the range as both NROH and SROH did not differ significantly among locations, groups (southeast vs. northwest), or areas (Vancouver Island, mainland coast of BC, Haida Gwaii, and SE Alaska; ANOVA, p > 0.05; Supplementary Figure 5). Furthermore, ROH sharing across locations did not show a geographical pattern either (Supplementary Figure 6). For example, location 8, on the central coast of BC, shared more ROHs with location 2 (southern Vancouver Island), and location 12 (Haida Gwaii) than any other neighbouring locations (Supplementary Figure 6). Estimated coalescent times of ROHs ranged from 48-500 generations before present. The two most widely shared ROHs, roh5_4 and roh5_6 on scaffold 5, had coalescent times of ∼400 and 227 generations before present, suggesting that Northern abalone may have gone through population contractions and expansions around those times.

**Figure 5.**
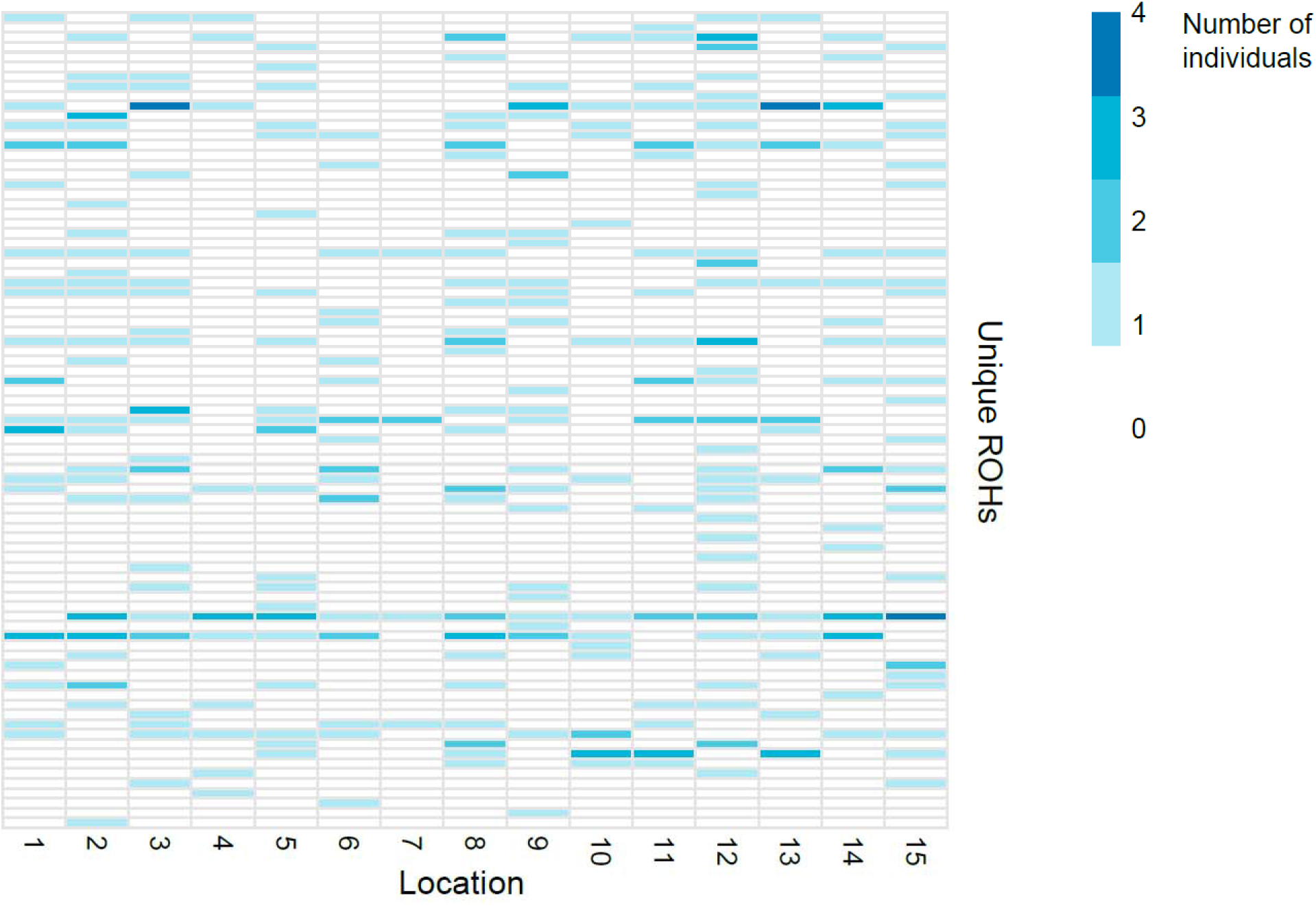
Distribution of ROHs across locations where each row represents a unique ROH, and the colour intensity represents how many individuals in a given location carry that unique ROH.

The analysis of rare alleles revealed that 15,761,026 variants (28% of all variants and 42% of rare variants) were private, i.e. present in one location only, and the vast majority of these (15,189,795 variants, 26% of all variants and 41% of rare variants) were present in only one individual, mostly in a heterozygous state (96.2%). These results highlight that a large portion of the total genomic variation in Northern abalone is made up of very rare alleles, that probably arose recently as new mutations following the bottleneck.

## Discussion

Using whole genome data and multiple analytical approaches, we have addressed fundamental questions related to the population structure and the population decline observed in Northern abalone from BC and SE Alaska, which will help inform their management in Canada and beyond. Our study shows that in Northern abalone, a species with high levels of genomic variation and weak differentiation, whole genome data are necessary to characterize population differentiation, genomic variation and demographic histories, and that reduced-representation approaches, including microsatellites, have failed to capture genomic signatures that may be relevant for management and conservation. While reduced-representation approaches present some advantages compared to whole genome sequencing, e.g. they are generally cheaper and require fewer genomics and bioinformatics resources, in the case of Northern abalone, a complete characterization of genomic differentiation could only be obtained by sampling the entirety of the genome.

We have also shown how informative data that are generally discarded can be, such as with rare alleles, or when used for a single specific application, such as ROHs. In this study, rare alleles were instrumental to better reconstruct population trends and characterize signals of population recovery and connectivity. Similarly, ROHs were used not only as indicators of current genetic inbreeding, but also to infer historical inbreeding and connectivity among distant sampling locations.

### Genomic differentiation in Northern abalone is concentrated and driven by selection and historical connectivity

Our analyses have corroborated previous genetic work reporting weak genome-wide differentiation (Withler et al., 2003; Dimond et al., 2024), but have also identified two distinguishable genetic groups, one including locations from Vancouver Island and the mainland coast of BC, and another including locations from Haida Gwaii and SE Alaska. Genomic differentiation was very concentrated in the genome, mostly on two scaffolds, 4 and 9, which explains why it had not been identified previously. In fact, we determined that a number of randomly sampled SNPs in the hundreds of thousands order of magnitude is required to detect population differentiation in Northern abalone, and that the population differentiation signal is quickly lost in analyses based on a lower number of random SNPs. In contrast, due to the concentrated nature of the genomic differentiation, a low number of highly differentiated SNPs, as little as 10, was sufficient to distinguish the two groups. Although low adult dispersal capabilities and relatively short pelagic larval phases would suggest stronger population structure than detected, in historically large populations like in the Northern abalone (*N_e_* = 300,000-500,000) genetic drift is weak and even minimal gene flow can further hamper differentiation (Slatkin, 1993). The combination of historically large population size, selection, weak genetic drift, and gene flow (current or historical), is likely what promotes the concentration of genomic differentiation in Northern abalone (Tigano & Friesen, 2016). Structural variants also likely enable the maintenance of these genomic patterns (Mérot et al., 2020), and their role in differentiation and local adaptation in Northern abalone is an open question that will be best addressed when a species-specific high quality reference genome becomes available.

These results generally align with other historically abundant abalone species, which tend to display overall weak to no population structure, with a few differentiated markers associated with latitudinal gradients (e.g., black abalone; Wooldridge et al., 2024), or environmental variation (e.g., red abalone and greenlip abalone; De Wit & Palumbi, 2013; Sandoval-Castillo et al., 2018).

In Northern abalone, the strongest signals of differentiation were mostly associated with genes involved in immune and stress response. Abalone worldwide are susceptible to disease. Withering syndrome, for example, almost exterminated black abalone throughout their range in California and Mexico (Rogers-Bennet et al. 2002). In laboratory exposure experiments, Northern abalone was the most susceptible to withering syndrome of all the Northeastern Pacific abalone species (Crosson & Friedman, 2018). Yet, diseases, including withering syndrome, have never been reported or tested as a major cause of population decline for Northern abalone. Differentiation at these immune and stress response genes may indicate that the two genetic groups may have been impacted differently by diseases in the past, leaving a stronger signature of selection, i.e. lower variation, in the southeast group. Two additional differentiated genes linked to reproductive functions (oocyte development [*PAQR5*] and sperm development [*SPAG16*]) curiously co-occurred in the same area of differentiation on scaffold 4. While these genes might hint at the evolution of reproductive isolation between the two genetic groups, this link is speculative in the absence of more information on phenotypic differentiation. Nonetheless, the stark contrast between negligible genome-wide differentiation and concentrated areas of differentiation suggests that either selection in these areas is strong, or that alternative alleles evolved in isolation, and were maintained by some recombination-suppression mechanism or selection against recombinants upon secondary contact (Guerrero & Hahn, 2017; Han et al., 2017; Nelson & Cresko, 2018).

Overall, the geographic patterns of population differentiation did not adhere to an isolation-by-distance model nor to expectations based on major oceanic current circulation in coastal BC, such as in the giant red sea cucumber (*Apostichopus californicus*), where population structure is consistent with the North Pacific Ocean current split into the Alaska Current northward and the California current southward off the coast of Vancouver Island (Xuereb et al., 2018). However, the population structure of the bat star (*Patiria miniata*) is more similar to the Northern abalone’s, with individuals from Haida Gwaii and SE Alaska forming a distinct group from the rest of BC, and a direct link to ocean circulation has been demonstrated (Sunday et al., 2014). Ocean circulation models of larval dispersal in the bat star explained these patterns of population structure well, showing how individuals from Vancouver Island were potentially connected with BC Central and North Coasts, and how isolated individuals from Alexander Archipelago in SE Alaska and, to a greater extent, Haida Gwaii were from the rest of the BC coast (Sunday et al. 2014). The different patterns of connectivity along the BC coast in different marine species may be explained by their different larval life histories, spawning behaviour, settlement cues, and habitat requirements. Species with longer pelagic larval phases are likely to be affected by broader current circulation and transported over longer distances than species with shorter larval phases, which are expected to be more strongly affected by finer-scale circulation patterns. However, pelagic larval duration may not necessarily be a good predictor of genetic distance (Esser et al. 2023) and other factors, unrelated to larval dispersal behaviour, such as deeper evolutionary histories, may leave a stronger signature than patterns of larval dispersal across a species range.

The greater historical *N_e_* and higher variation in the highly differentiated SNPs in the northwest group compared to the southeast group, together with the greatest mean individual heterozygosity in the Haida Gwaii region, is consistent with the hypothesis that Haida Gwaii, and potentially the Alexander Archipelago, may have been glacial refugia during the Last Glacial Maximum for many marine and terrestrial species (Shafer et al., 2010), including the Northern abalone. Furthermore, the sharing of fixed differences between the southeast group and the two northernmost locations on Haida Gwaii and SE Alaska further support that these two northernmost locations may have been historically connected with Northern abalone from the mainland coast of BC and/or represent areas of secondary contact following the Last Glacial Maximum, but independent in space or time from each other. Taken together, these results indicate that genomic differentiation in the northern range of the Northern abalone is primarily driven by historical segregation and selection in a few concentrated areas of the genome.

### Genomic analyses reveal a strong genomic signature of population decline but no inbreeding depression

Despite high overall genomic variation, we identified bottlenecks in both genetic groups resulting in 98-99% effective population reductions, with steep declines over approximately six generations, that were not detected in previous genetic analyses (Withler et al. 2003; Dimond et al. 2024). The population trends reconstructed here from genomic data are consistent with the evidence of a severe decline inferred from various sources of knowledge, including zooarcheological, historical, traditional and western science evidence (Lee et al., 2019): Northern abalone had large effective population sizes, in the northwest more than in the southeast, and in only a few generations, earlier in the southeast, numbers plummeted dramatically. However, it is difficult to temporally calibrate our demographic reconstructions due to unknown recombination rates, varying greatly within molluscs (Stapley et al., 2017), and uncertainty associated with generation time. In fact, our reconstructions based on three different recombination rates illustrate the effect of varying recombination estimates on the dating of demographic events, which becomes unattainable if accurate recombination rates are not available for the species of interest. Additionally, generation time for Northern abalone may vary between 2-10 years and be affected by variation in sea otter predation, food availability, and growth rates across the species range (Obradovich et al. 2021), thus adding another layer of uncertainty in the dating of demographic events.

Our analyses hint at the start of a timid recovery in the last 8-10 generations, possibly a results of the closure of all Northern abalone harvest in BC in 1990. The signal of recovery from the demographic reconstructions, based on patterns of LD, is supported by a pervasive excess of rare alleles across the genome, which cannot be attributed to selection, but rather indicate population expansion. Further, the high proportion of singletons/doubletons – more than 1 in 4 – indicate that these alleles are likely new mutations that have arisen recently, and have not had the time to increase in frequency due to drift or selection. The abundance of these rare alleles was thought to be the result of high mutation rates in abalone species (De Wit & Palumbi, 2013), but a recent study shows that white abalone (*Haliotis sorenseni*) has mutation rates comparable to vertebrates with similar longevity and generation times (Wooldridge et al., 2025), suggesting that population recovery is occurring over longer timescales. Indications of population recovery from the genomic data are supported by empirical observations of increasing densities of Northern abalone, though mostly of small juveniles (Lee et al., 2019; Obradovich et al., 2021). These results highlight how informative rare variants, which are generally discarded from population genomics analyses, can be about recent demography. Had we not considered the allele frequency of the numerous SNPs identified, the amount of variation alone would have suggested good genetic health, as concluded previously based on the analysis of fewer markers (Withler et al., 2003; Dimond et al., 2024).

Despite the dramatic population decline, we did not identify an amount nor size distribution of ROHs indicative of inbreeding depression. Large population sizes and existing high genomic variation before the bottleneck may have prevented detectable inbreeding depression, or the abundance of new mutations that occurred after the bottleneck may have masked the lack of genomic variation that characterizes ROHs. As new mutations tend to be deleterious (Ohta, 1973), such an abundance of rare alleles could itself pose a risk to the genetic health of recovering Northern abalone. Increasing genetic load (mutation load), compounded by the issue that population size is drastically smaller than before the bottleneck and new deleterious mutations may spread more quickly, may further slow the species recovery (e.g., drift load; Bertorelle et al., 2022). In the Northern elephant seal, for example, a severe bottleneck reducing the species to a few individuals did not seem to result in inbreeding depression, but the genetic load due to the accumulation of new mutations during the population expansion that followed appeared to have slowed population recovery (Hoffman et al., 2024).

ROHs are identical by descent segments and as such can be used as genetic markers to infer not only inbreeding depression, but also recent demography and connectivity. One advantage of ROHs over SNPs is that they are less susceptible to homoplasy, i.e. when the same variant arises independently in different individuals and locations. As such, they are more reliable to infer connectivity. In our dataset, homoplasy would be an extremely unlikely explanation for the sharing of one or both breakpoints of hundreds-of-kb-long ROHs, often of the same lengths, across individuals. In our dataset, sharing of ROHs across distant sampling locations suggests that periods of localized inbreeding in the past were followed by dispersal and gene flow, probably associated with range expansions and contractions, which carried those ROHs across long distances. During the Holocene only, the coast of BC and SE Alaska was highly dynamic, with sea levels much higher or lower than today, and changing rapidly (Josenhans et al., 1995). Changes in coastline, exposed land, and currents might have increased or decreased connectivity over time across the Northern abalone range, potentially explaining the spatial distribution of ROHs and blocks of differentiation in BC and SE Alaska.

### Management implications

Northern abalone is managed as a single Designatable Unit in Canada (Species at Risk Public Registry, 2025). Here, we have found evidence of population differentiation, albeit concentrated in a few genomic areas, between two geographically broad genetic groups. The concentrated nature of population differentiation raises several questions for the management of Northern abalone: how pervasive does the genome differentiation need to be for two groups of individuals to be considered distinct and thus managed as such? And, do those differentiated genomic regions have phenotypic and/or adaptive significance? An increasing number of studies are showing that differentiation can be extremely concentrated in the genome even among what are considered reproductively isolated species (Campagna et al., 2017; Toews et al., 2016), and in many cases a link between these genomic regions and the factors leading to reproductive or ecological isolation between taxa, including populations, ecotypes, or species, has been demonstrated (Barrett et al., 2019; Loveland et al., 2025; Turbek et al., 2021). In Northern abalone, differentiation was mostly associated with genes related to immune response and reproductive functions, but their relative fitness effects are unknown. Understanding if and how these genes affect phenotypes and fitness would require functional validation of different variants or haplotypes and assessment of their fitness effect, but this research is hampered by their relatively long generation times, poor genomic resources for the species (and molluscs more broadly), and their conservation status limiting the necessary experiments. This said, our analyses show that, despite overall low genomic differentiation, the Northern abalone do not represent a single panmictic demographic unit, and that current gene flow among our sampling locations is lower than previously argued (Withler et al., 2003; Dimond et al., 2024). Finally, our demographic reconstructions support existing evidence of a severe population decline and a timid population recovery, at least across the northern part of the Northern abalone range (BC and SE Alaska), which suggests that fishery closures have been effective in slowing the decline and promoting recovery. However, the examined locations are still far from pre-bottleneck abundances, and continued conservation efforts are crucial to enable further recovery and the long-term persistence of the species.

## Acknowledgements

We would like to thank all the First Nation and Fisheries and Oceans (DFO) biologists who contributed to sampling underlying this work, including specifically the following who led trips where Northern abalone DNA samples were collected: Barbara Lucas, Joel Harding, Joanne Lessard, Seaton Taylor. The Haida Nation (Dan McNeill) contributed some of the samples from Haida Gwaii. The Kitasoo Xai Xai’s Nation (Sandie Hankewich) provided some of the samples from the Central Coast. We thank Genome Quebec for whole genome library preparation and sequencing, and Genome Quebec for library preparation and sequencing. Computational analyses were conducted on the Shared Services Canada General Purpose Science Cluster with support from the Government of Canada through the Genomics Research and Development Initiative – Genomic Adaption and Resilience to Climate Change (GenARCC) project (2022-2027). We thank Braden Judson, Tim Healy, and Kyle Wellband for feedback and helpful discussions that helped improve the manuscript.

## Author Contributions

AT performed all analyses and wrote the manuscript. KL performed DNA extractions.

JS and EBR selected samples and locations to include in the study. DC coordinated sampling permits and sample collection by DFO and First Nations. SCH and MDM helped develop the connectivity component of the study. AT, ECH, EBR, and DB conceived the study.

DB acquired funds to support the study. All authors provided critical feedback on analyses and the manuscript.

## Data availability

Whole genome resequencing data will be submitted to ENA and scripts accompanying analysis will be public on GitHub/Zenodo.

---

**Supplementary Figure 1.**
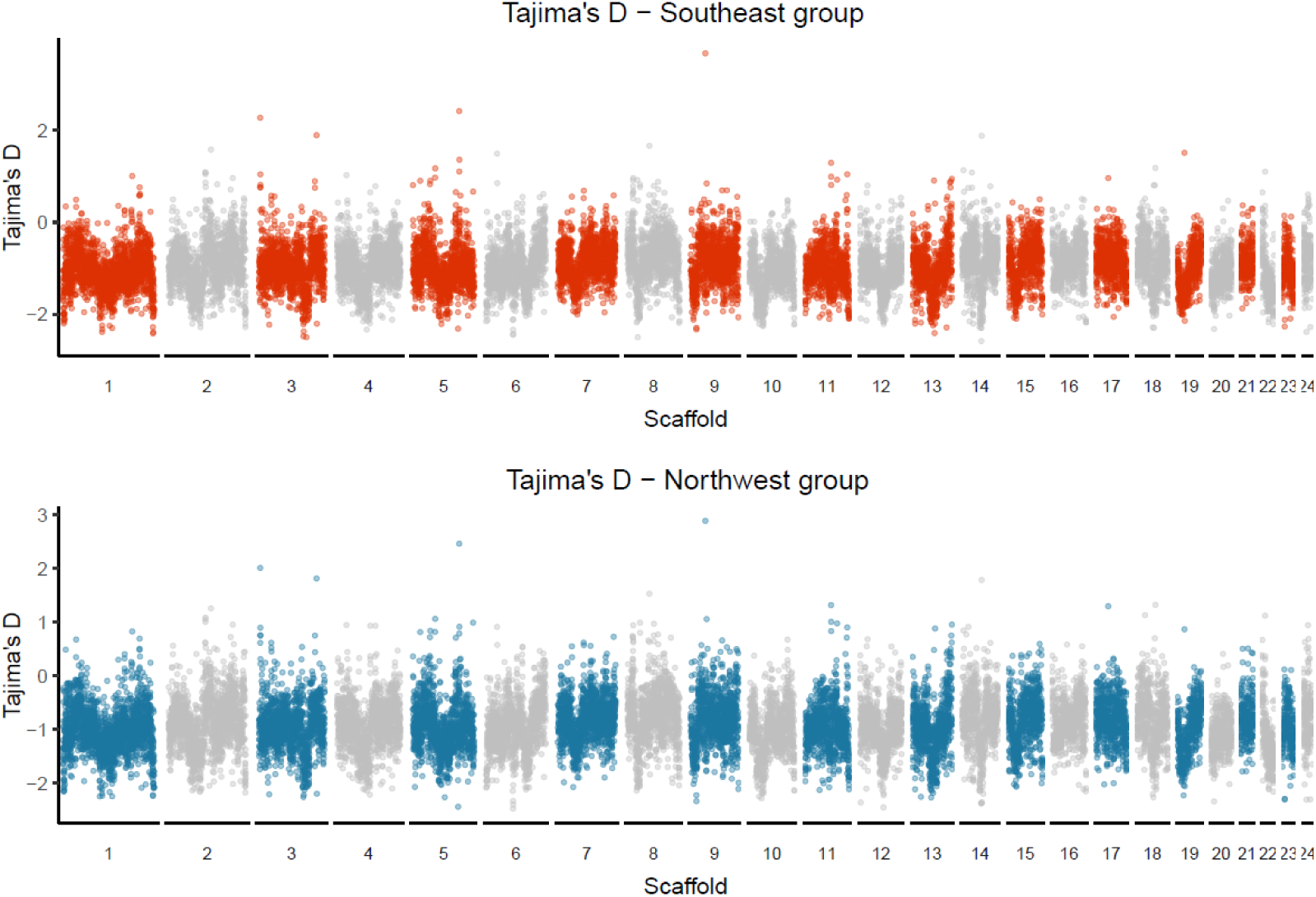
Manhattan plots of Tajima’s D values estimated in 50 kb moving windows in the southeast group (upper panel) and in the northwest group (lower panel).

**Supplementary Figure 2.**
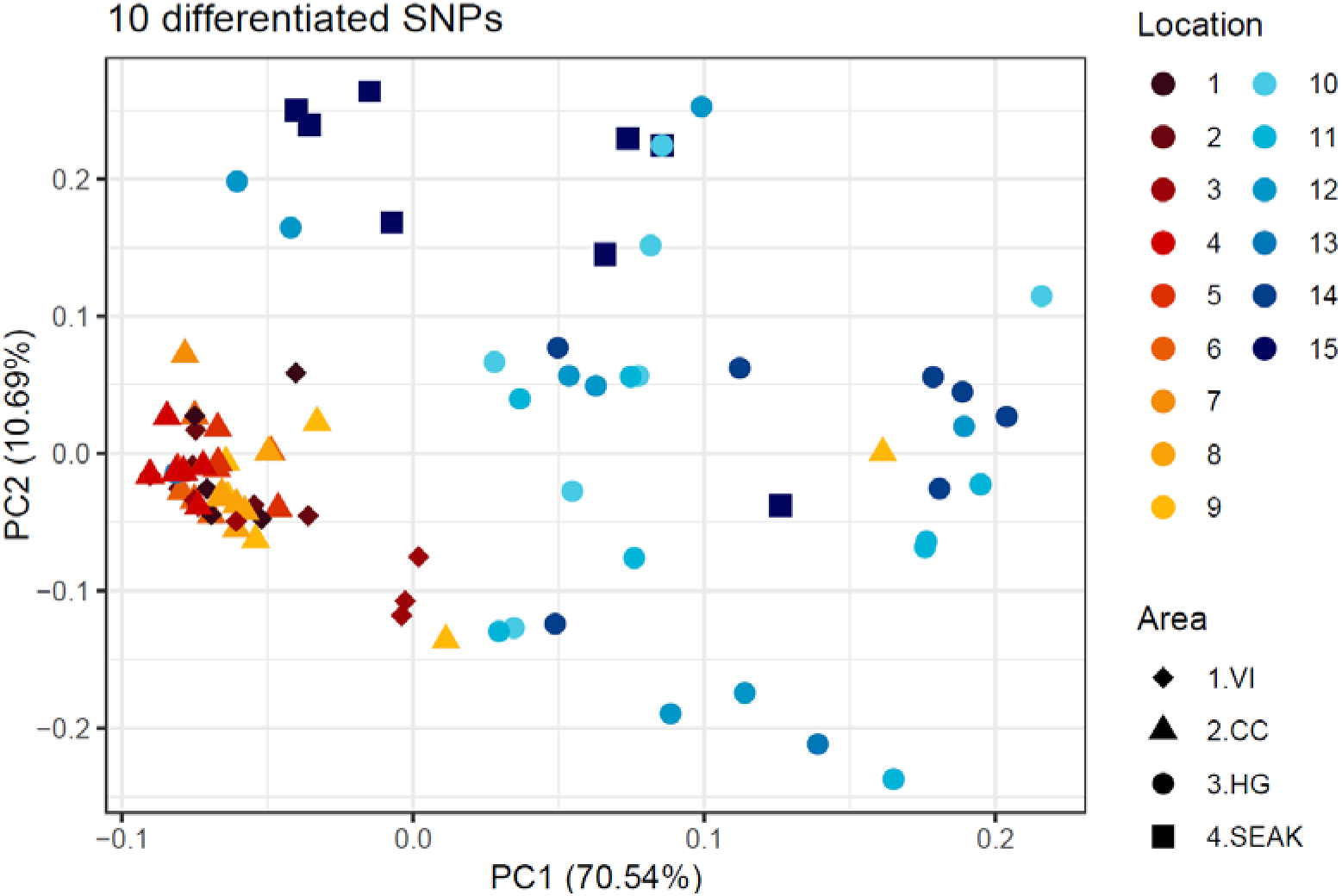
Plot of PCA based on 10 highly differentiated SNPs. VI = Vancouver Island, CC = Central Coast, HG = Haida Gwaii, and SEAK = Southeastern Alaska.

**Supplementary Figure 3.**
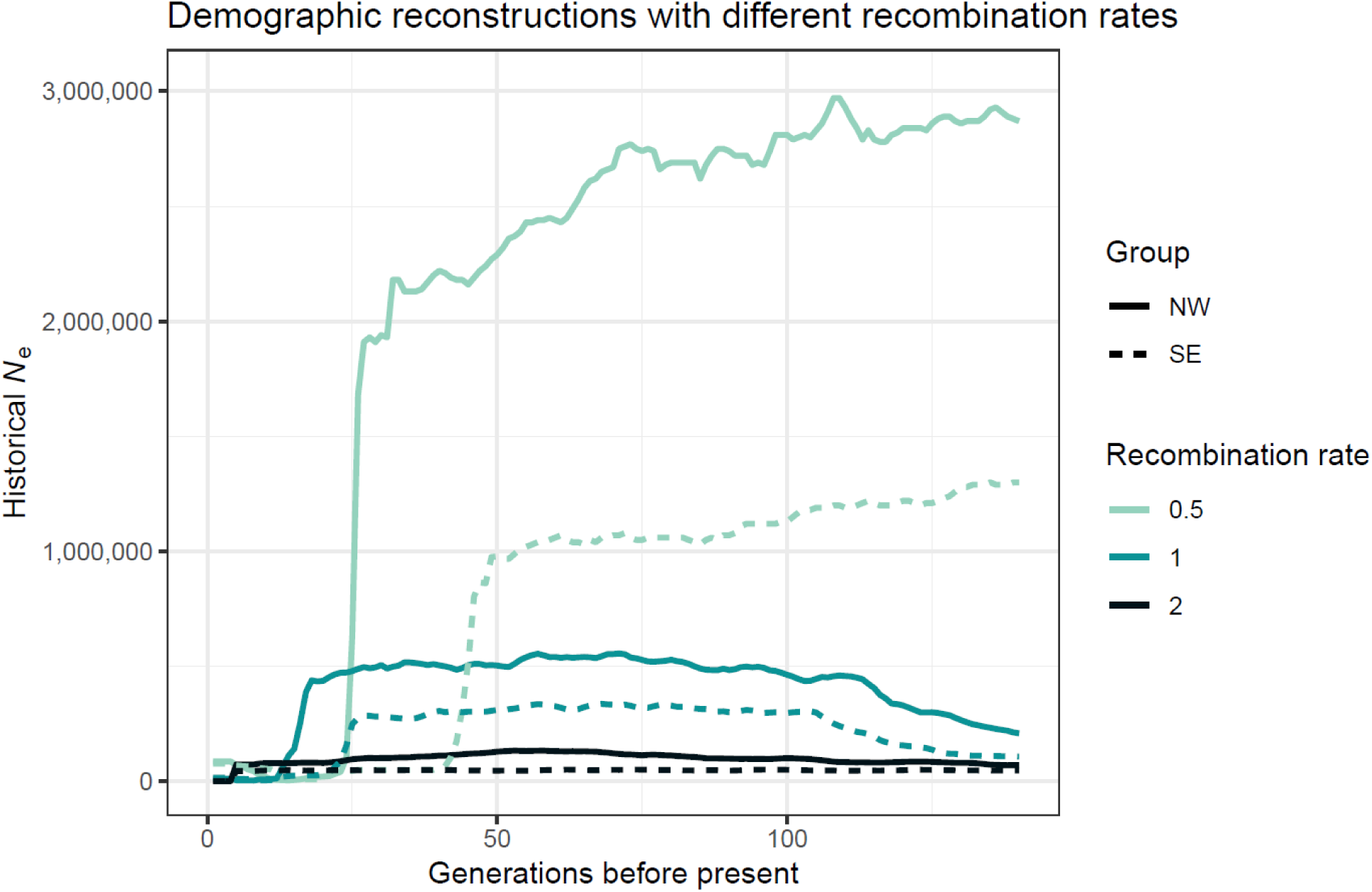
Plot summarizing demographic reconstructions for each genetic group (NW = northwest, SE = southeast) using three different recombination rates.

**Supplementary Figure 4.**
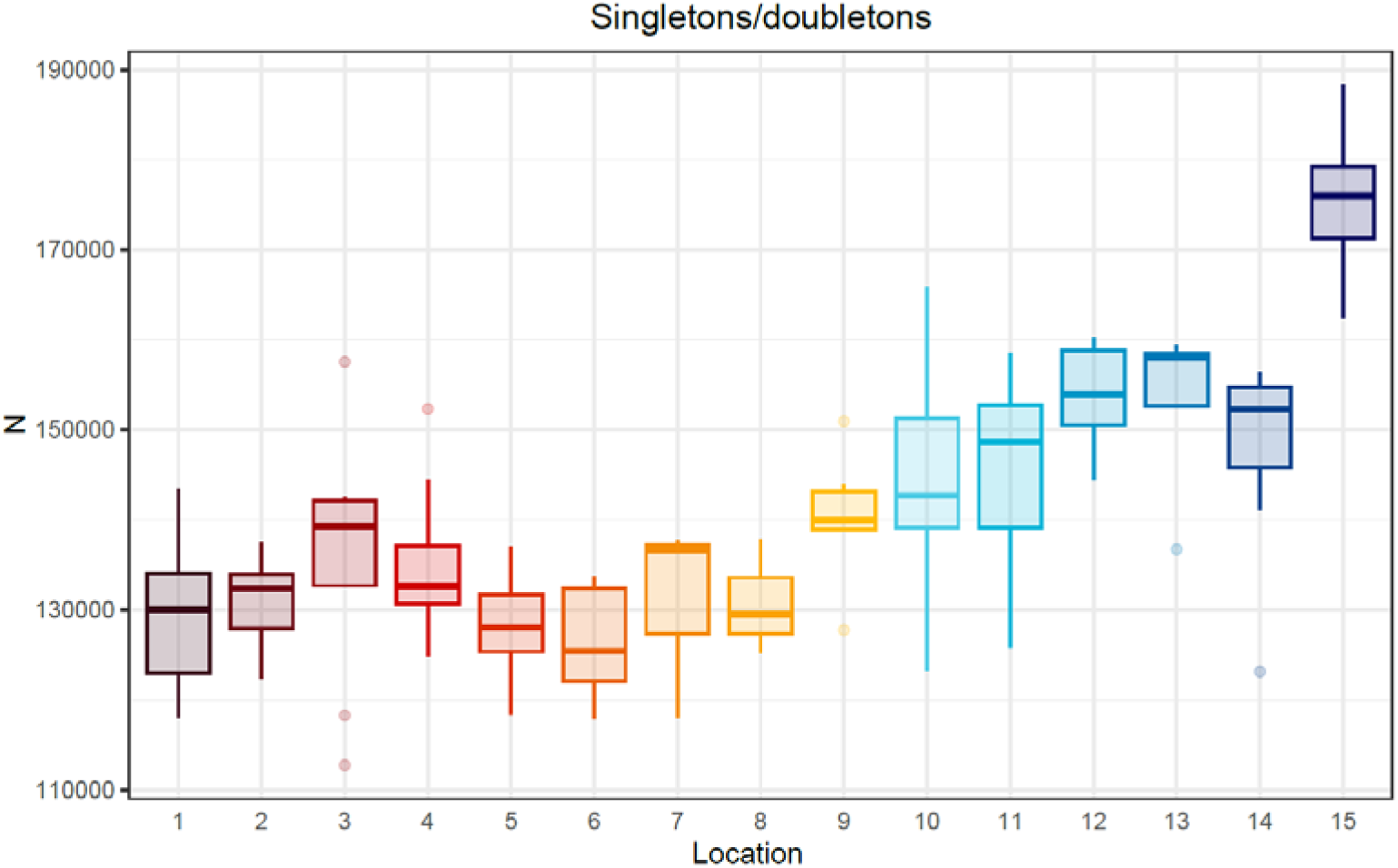
Boxplot showing the distribution of singletons/doubletons, variants occurring in only one individual as either homozygous or heterozygous, across sampling locations.

**Supplementary Figure 5.**
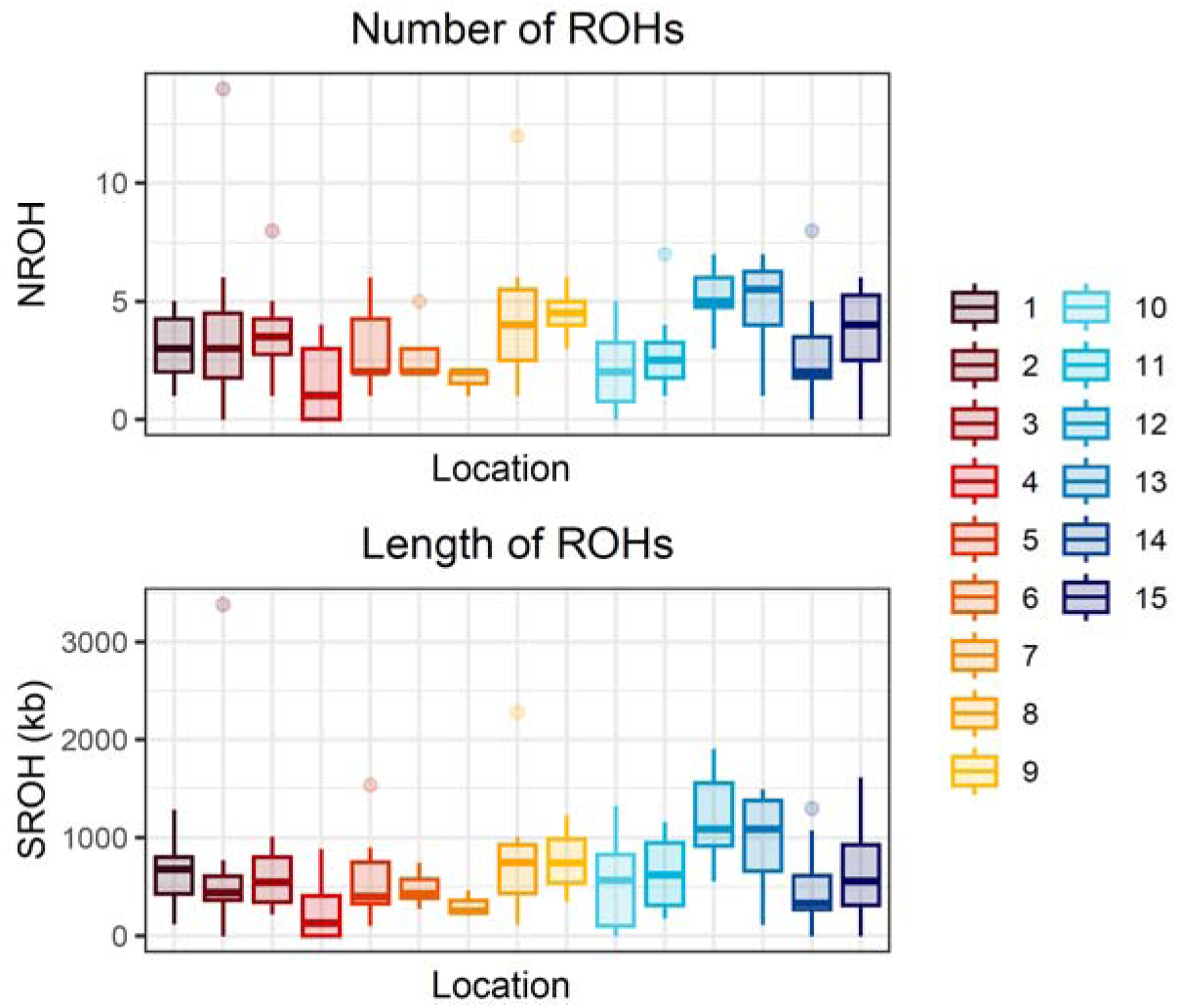
Boxplots showing the distribution of number (NROH) and length (SROH) of ROHs across sampling locations.

**Supplementary Figure 6.**
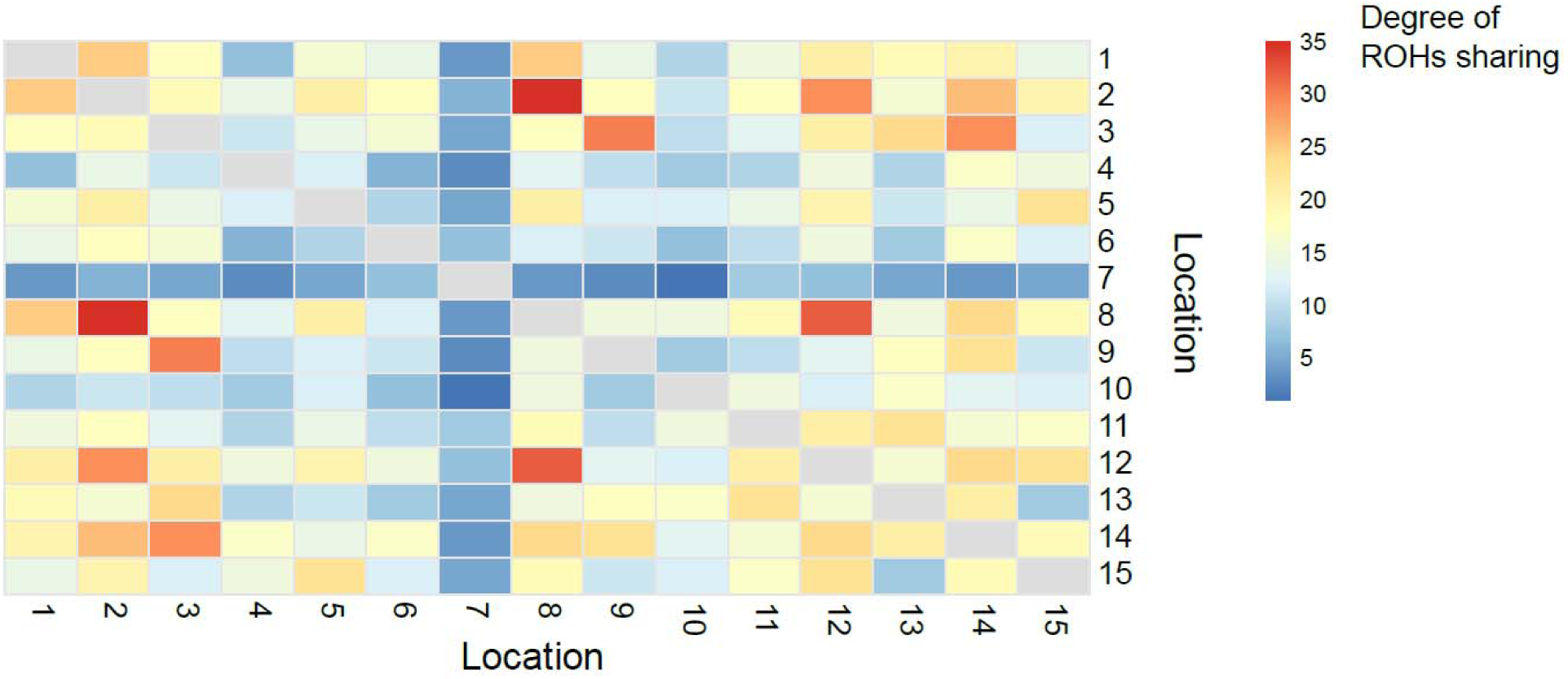
Heatmap showing the degree of overlap of ROHs from different sampling locations.

**Supplementary Table 1.**
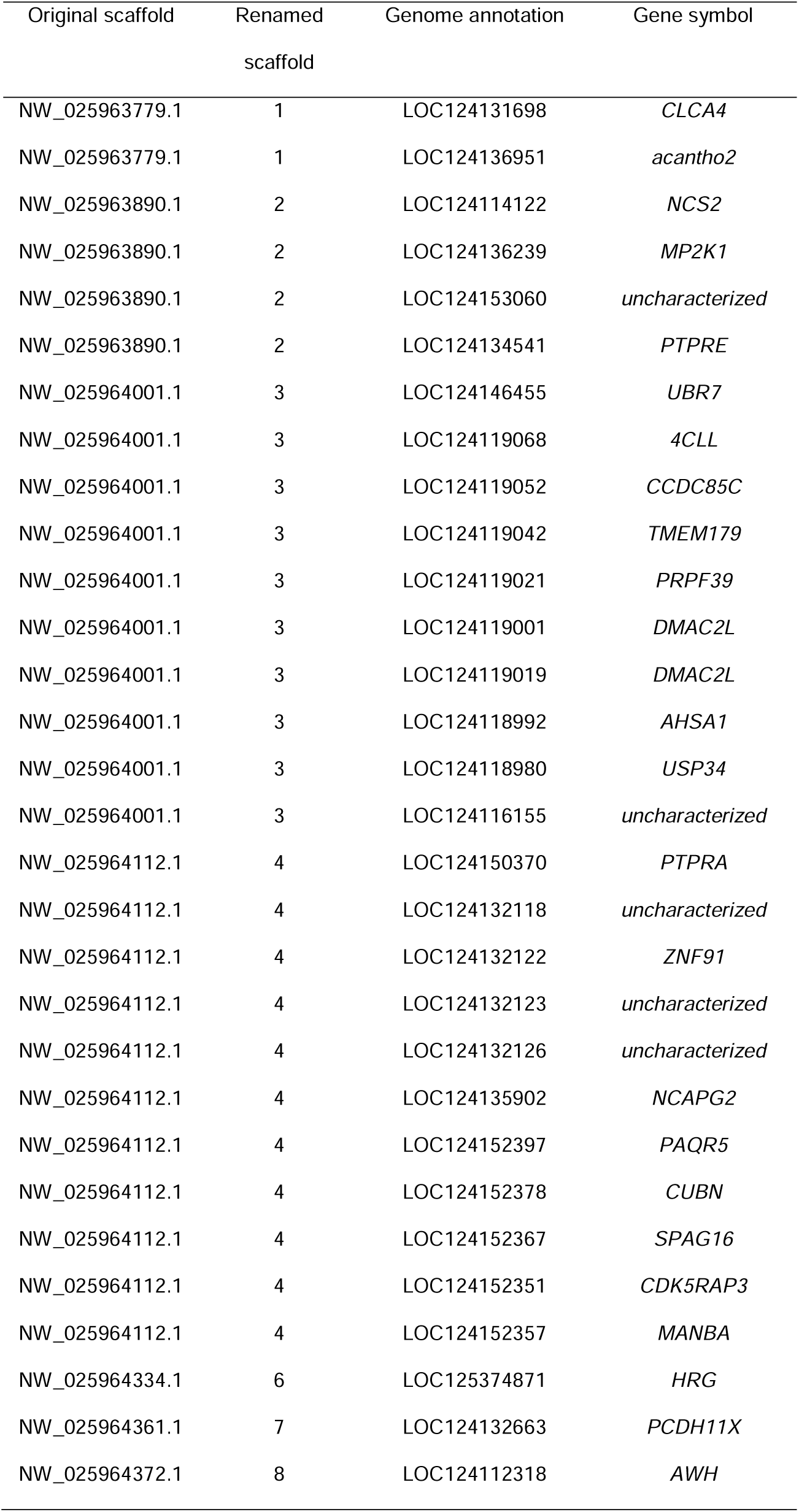
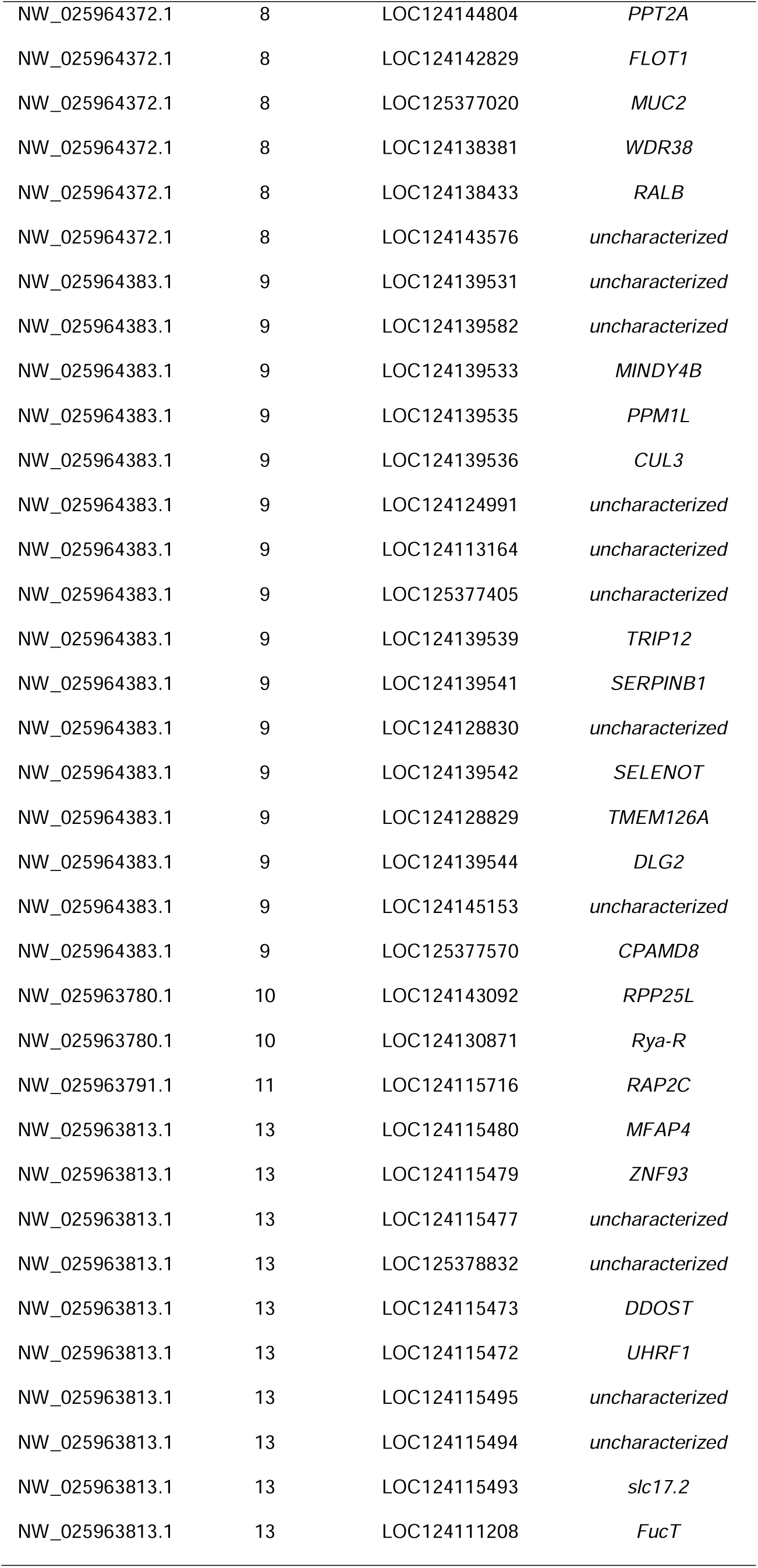
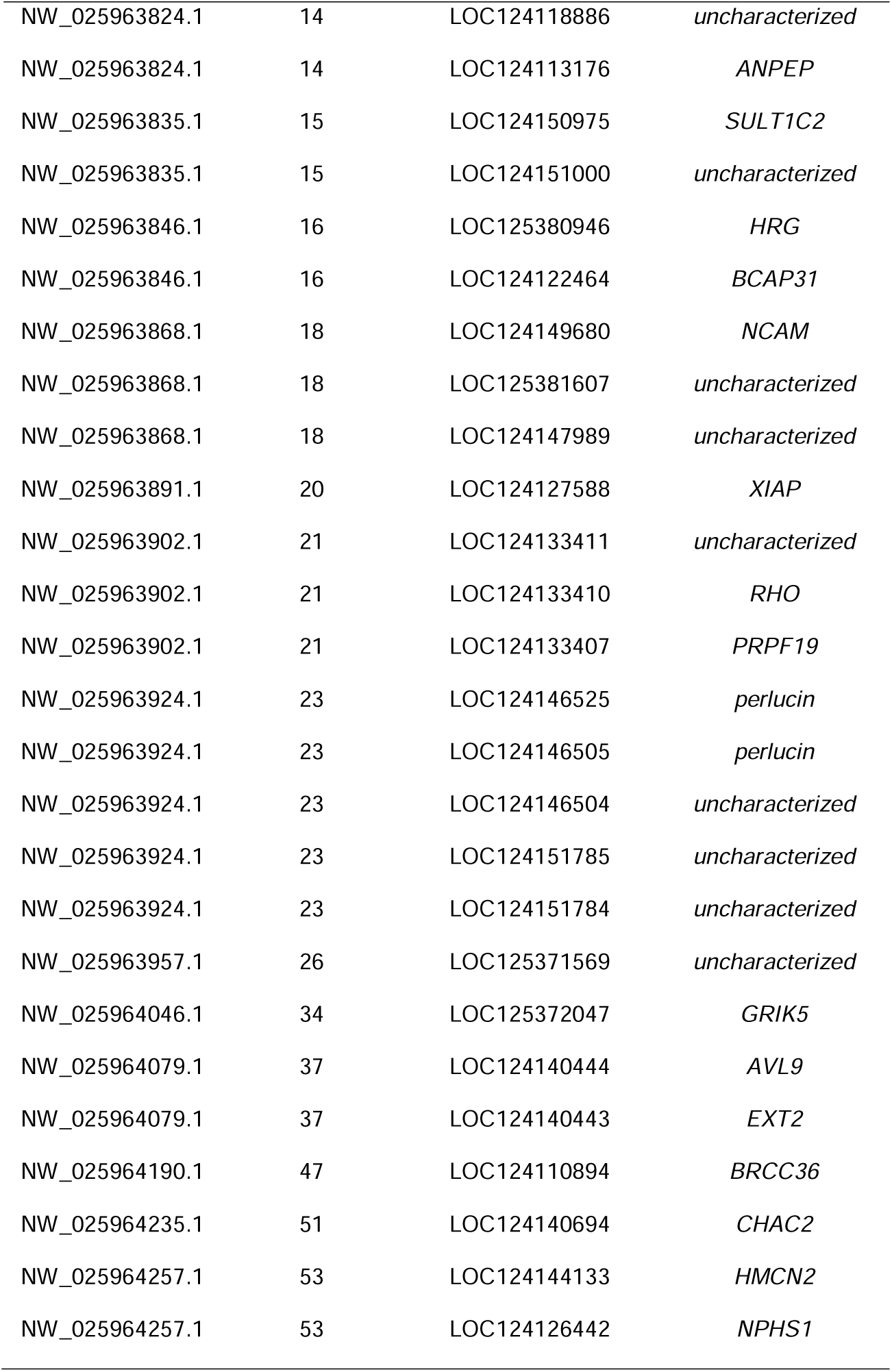
List of genes affected by the 1,000 most differentiated SNPs.

| Original scaffold | Renamed scaffold | Genome annotation | Gene symbol |
| --- | --- | --- | --- |
| NW_025963779.1 | 1 | LOC124131698 | *CLCA4* |
| NW_025963779.1 | 1 | LOC124136951 | *acantho2* |
| NW_025963890.1 | 2 | LOC124114122 | *NCS2* |
| NW_025963890.1 | 2 | LOC124136239 | *MP2K1* |
| NW_025963890.1 | 2 | LOC124153060 | *uncharacterized* |
| NW_025963890.1 | 2 | LOC124134541 | *PTPRE* |
| NW_025964001.1 | 3 | LOC124146455 | *UBR7* |
| NW_025964001.1 | 3 | LOC124119068 | *4CLL* |
| NW_025964001.1 | 3 | LOC124119052 | *CCDC85C* |
| NW_025964001.1 | 3 | LOC124119042 | *TMEM179* |
| NW_025964001.1 | 3 | LOC124119021 | *PRPF39* |
| NW_025964001.1 | 3 | LOC124119001 | *DMAC2L* |
| NW_025964001.1 | 3 | LOC124119019 | *DMAC2L* |
| NW_025964001.1 | 3 | LOC124118992 | *AHSA1* |
| NW_025964001.1 | 3 | LOC124118980 | *USP34* |
| NW_025964001.1 | 3 | LOC124116155 | *uncharacterized* |
| NW_025964112.1 | 4 | LOC124150370 | *PTPRA* |
| NW_025964112.1 | 4 | LOC124132118 | *uncharacterized* |
| NW_025964112.1 | 4 | LOC124132122 | *ZNF91* |
| NW_025964112.1 | 4 | LOC124132123 | *uncharacterized* |
| NW_025964112.1 | 4 | LOC124132126 | *uncharacterized* |
| NW_025964112.1 | 4 | LOC124135902 | *NCAPG2* |
| NW_025964112.1 | 4 | LOC124152397 | *PAQR5* |
| NW_025964112.1 | 4 | LOC124152378 | *CUBN* |
| NW_025964112.1 | 4 | LOC124152367 | *SPAG16* |
| NW_025964112.1 | 4 | LOC124152351 | *CDK5RAP3* |
| NW_025964112.1 | 4 | LOC124152357 | *MANBA* |
| NW_025964334.1 | 6 | LOC125374871 | *HRG* |
| NW_025964361.1 | 7 | LOC124132663 | *PCDH11X* |
| NW_025964372.1 | 8 | LOC124112318 | *AWH* |
| NW_025964372.1 | 8 | LOC124144804 | *PPT2A* |
| NW_025964372.1 | 8 | LOC124142829 | *FLOT1* |
| NW_025964372.1 | 8 | LOC125377020 | *MUC2* |
| NW_025964372.1 | 8 | LOC124138381 | *WDR38* |
| NW_025964372.1 | 8 | LOC124138433 | *RALB* |
| NW_025964372.1 | 8 | LOC124143576 | *uncharacterized* |
| NW_025964383.1 | 9 | LOC124139531 | *uncharacterized* |
| NW_025964383.1 | 9 | LOC124139582 | *uncharacterized* |
| NW_025964383.1 | 9 | LOC124139533 | *MINDY4B* |
| NW_025964383.1 | 9 | LOC124139535 | *PPM1L* |
| NW_025964383.1 | 9 | LOC124139536 | *CUL3* |
| NW_025964383.1 | 9 | LOC124124991 | *uncharacterized* |
| NW_025964383.1 | 9 | LOC124113164 | *uncharacterized* |
| NW_025964383.1 | 9 | LOC125377405 | *uncharacterized* |
| NW_025964383.1 | 9 | LOC124139539 | *TRIP12* |
| NW_025964383.1 | 9 | LOC124139541 | *SERPINB1* |
| NW_025964383.1 | 9 | LOC124128830 | *uncharacterized* |
| NW_025964383.1 | 9 | LOC124139542 | *SELENOT* |
| NW_025964383.1 | 9 | LOC124128829 | *TMEM126A* |
| NW_025964383.1 | 9 | LOC124139544 | *DLG2* |
| NW_025964383.1 | 9 | LOC124145153 | *uncharacterized* |
| NW_025964383.1 | 9 | LOC125377570 | *CPAMD8* |
| NW_025963780.1 | 10 | LOC124143092 | *RPP25L* |
| NW_025963780.1 | 10 | LOC124130871 | *Rya-R* |
| NW_025963791.1 | 11 | LOC124115716 | *RAP2C* |
| NW_025963813.1 | 13 | LOC124115480 | *MFAP4* |
| NW_025963813.1 | 13 | LOC124115479 | *ZNF93* |
| NW_025963813.1 | 13 | LOC124115477 | *uncharacterized* |
| NW_025963813.1 | 13 | LOC125378832 | *uncharacterized* |
| NW_025963813.1 | 13 | LOC124115473 | *DDOST* |
| NW_025963813.1 | 13 | LOC124115472 | *UHRF1* |
| NW_025963813.1 | 13 | LOC124115495 | *uncharacterized* |
| NW_025963813.1 | 13 | LOC124115494 | *uncharacterized* |
| NW_025963813.1 | 13 | LOC124115493 | *slc17.2* |
| NW_025963813.1 | 13 | LOC124111208 | *FucT* |
| NW_025963824.1 | 14 | LOC124118886 | *uncharacterized* |
| NW_025963824.1 | 14 | LOC124113176 | *ANPEP* |
| NW_025963835.1 | 15 | LOC124150975 | *SULT1C2* |
| NW_025963835.1 | 15 | LOC124151000 | *uncharacterized* |
| NW_025963846.1 | 16 | LOC125380946 | *HRG* |
| NW_025963846.1 | 16 | LOC124122464 | *BCAP31* |
| NW_025963868.1 | 18 | LOC124149680 | *NCAM* |
| NW_025963868.1 | 18 | LOC125381607 | *uncharacterized* |
| NW_025963868.1 | 18 | LOC124147989 | *uncharacterized* |
| NW_025963891.1 | 20 | LOC124127588 | *XIAP* |
| NW_025963902.1 | 21 | LOC124133411 | *uncharacterized* |
| NW_025963902.1 | 21 | LOC124133410 | *RHO* |
| NW_025963902.1 | 21 | LOC124133407 | *PRPF19* |
| NW_025963924.1 | 23 | LOC124146525 | *perlucin* |
| NW_025963924.1 | 23 | LOC124146505 | *perlucin* |
| NW_025963924.1 | 23 | LOC124146504 | *uncharacterized* |
| NW_025963924.1 | 23 | LOC124151785 | *uncharacterized* |
| NW_025963924.1 | 23 | LOC124151784 | *uncharacterized* |
| NW_025963957.1 | 26 | LOC125371569 | *uncharacterized* |
| NW_025964046.1 | 34 | LOC125372047 | *GRIK5* |
| NW_025964079.1 | 37 | LOC124140444 | *AVL9* |
| NW_025964079.1 | 37 | LOC124140443 | *EXT2* |
| NW_025964190.1 | 47 | LOC124110894 | *BRCC36* |
| NW_025964235.1 | 51 | LOC124140694 | *CHAC2* |
| NW_025964257.1 | 53 | LOC124144133 | *HMCN2* |
| NW_025964257.1 | 53 | LOC124126442 | *NPHS1* |

